# Surface modification of PDMS-based microfluidic devices with collagen using polydopamine as a spacer to enhance primary human bronchial epithelial cell adhesion

**DOI:** 10.1101/2020.11.09.375709

**Authors:** Mohammadhossein Dabaghi, Shadi Shahriari, Neda Saraei, Kevin Da, Abiram Chandiramohan, Ponnambalam Ravi Selvaganapathy, Jeremy A. Hirota

**Author notes:** Corresponding author: Jeremy A. Hirota.

## Abstract

Polydimethylsiloxane (PDMS) is a silicone-based synthetic material that is used in various biomedical applications due to its properties, including transparency, flexibility, permeability to gases, and ease of use. Though PDMS facilitates and realizes the fabrication of complicated geometries at the micro and nano scales, it does not optimally interact with cells for adherence and proliferation. Different strategies have been proposed to render PDMS to enhance cell attachment. The majority of these surface modification techniques have been offered for a static cell culture system. However, dynamic cell culture systems such as organ-on-a-chip devices are demanding platforms that recapitulate the complexity of a living tissue microenvironment. For organ-on-a-chip platforms, PDMS surfaces are usually coated by ECM proteins, which occur as a result of physical, weak bonding between PDMS and ECM proteins, and this binding can be degraded when it is exposed to shear stresses. This work reports static and dynamic coating methods to covalently bind collagen within a PDMS-based microfluidic device using polydopamine (PDA). These coating methods were evaluated using water contact angle measurement and atomic force microscopy (AFM) to find the optimum coating conditions. The biocompatibility of collagen-coated PDMS devices was assessed by culturing primary human bronchial epithelial cells (HBECs) in microfluidic devices. It was shown that both PDA coating methods could be used to bind collagen, thereby improving cell adhesion (around three times higher) without showing any discernible difference. These results suggested that such a surface modification can be used to coat an extracellular matrix protein onto PDMS-based microfluidic devices.

## Introduction

The miniaturization of biomedical devices has been increasing in demand, leading to new fabrication approaches to produce such devices that can be used in various diagnostic and biological applications^1^. The use of PDMS has been evident in a variety of biomedical applications as biomedical devices due to biocompatibility, low barriers to cost, and fabrication, especially for microfluidic applications where rapid prototyping and inexpensive prototyping can allow for long term usage^2^. PDMS possesses these beneficial properties for applications ranging from cell sorting to organ-on-a-chip devices. Organ-on-a-chip devices have attracted wide attention from pharmaceutical companies and academic laboratories due to the potential of these devices to emulate physiological phenomena at micro-scale^3,4^. Organ-on-a-chip platforms can recapitulate the complexity of tissues and organs to some extent by combining cellular and extracellular cues in a chip. In addition to promising features such as providing tissue barriers and hydrodynamic forces that organ-on-a-chip devices offer, the inner surface of organ-on-a-chip devices can be coated by extracellular matrix (ECM) components to resemble the native cellular microenvironment and improve cellular adhesion ^3^.

The majority of organ-on-a-chip devices have been made of PDMS because of the unique characteristics of PDMS. However, the PDMS surface of microfluidic devices needs to be tailored prior to the cell culture so that cells can uniformly adhere to the surface, grow, and proliferate. Although a conventional surface modification technique such as oxygen plasma^5^ can be used to introduce hydroxyl groups onto PDMS surface, thereby improving the cell attachment, such a modification is not stable due to hydrophobic recovery of PDMS, which requires cell culture immediately following plasma treatment to obtain optimal cell attachment. Another approach is to coat an ECM-based component such as collagen or fibronectin by physical adsorption that occurs as the result of weak electrostatic and van der Waals interactions between the ECM-based material and PDMS. This method has been followed since the earlier works in the field of microfluidics and organ-on-a-chip to improve cell adhesion^6,7^. Since the attachment between the ECM-based component and PDMS is physical, it may degrade over time and compromise the homogeneity of cultured cells. A more favorable approach is to covalently bond ECM proteins to PDMS using a spacer or linker such as (3-aminopropyl)triethoxy silane (APTES)^8^. The use of silane-based materials has two disadvantages: First, it requires an oxygen plasma treatment to introduce hydroxyl groups onto PDMS prior to silane coating, which may not be accessible to every lab. Second, silane molecules are toxic and may cause cell death if some regions are not fully covered by ECM proteins.

To capture the benefit of ECM proteins coating while addressing the shortcomings of other surface modification techniques to graft an ECM protein or a combination of them on PDMS, a simple method is used to coat polydopamine (PDA) onto PDMS as biomolecule agent. Dopamine (DA) in an alkaline solution (pH = 8.5) undergoes an oxidative polymerization and forms PDA layers on submerged surfaces ^9^. PDA itself ^10^ or as a coating agent ^11^ can be utilized to improve the cell adhesion on PDMS. In fact, PDA, as a linker, can interact with the amine groups of ECM proteins and covalently bind them onto the surface ^12^. Though PDA coating has been widely used to render various substrates ^12–14^ including PDMS ^10,15,16^ for improving cell attachment, it has not been used to coat an ECM-based protein in a microfluidic device. In a recent study, it has been demonstrated that PDA coating inside a microfluidic device is stable enough to be stored for months at room temperature or can be sterilized under extreme conditions without any negative impact on cell viability and functionality^16^. Although it was shown that PDA coating would enhance the cell attachment, PDA cannot communicate and interact with cells in the way that ECM-based materials may do.

## Methods

### Device fabrication

PDMS microfluidic devices were producing using Sylgard^®^ 184 silicone elastomer kit mixing at a 10:1 ratio of base and cross-linker agent. It was degassed for about 30 minutes and then cast onto a silicon wafer mold with two silicone tubes as the inlet and outlet to cure for at least an hour on a hot plate or in an oven at 85 °C. After curing, the casted PDMS devices were removed where they were then oxygen-plasma or flame ^17^ treated with another layer of PDMS (oxygen plasma treatment was performed for two minutes at a pressure of 900 mTorr in a Harrick Plasma cleaner machine.) After exposure, contact between the channel layer and the blank PDMS layer occurred where bonding between silanol groups after oxygen-plasma or flame oxidation exposure could occur to seal the two layers. The device could then be set to cure on a hot plate or in an oven at 85 °C for 24 hours.

### Polydopamine coating

Dopamine hydrochloride was produced from Sigma Aldrich and prepared at a 1, 2, and 5 mg mL^−1^ concentration into an 8.5 pH phosphate-based-buffer (PBS). Upon stirring, devices to be coated were connected in series with Tygon tubing. PDA solution was run through devices at various flow rates (0.5, 1, 3, 6, and 9 mL min^−1^) with a peristaltic pump for 24 or 48 hours. For static PDA coating, the microfluidic devices were filled with the DA solution with the desired concentration for 24 or 48 hours. After PDA coating, the devices were rinsed with PBS (pH = 7.4) and DI water.

### Contact angle measurement and AFM

After PDA coating and rinsing the microfluidic devices, they were cut along the length of the channel to expose the inner coated surfaces to the atmosphere. Then, the samples were dried at room temperature overnight prior to surface analysis measurements. For contact angle measurements, a 2-μL water droplet was used.

For AFM, we used a silicon nitride cantilever that had a spring constant of 0.4 N m, and it was adjusted to automatically scan the treated and non-treated surfaces at a scan rate of 1 Hz by acquiring 512 samples per line. The instrument software (NanoScope Analysis) was acquired for image analysis.

### Cell culture and Calcein AM imaging

#### Human ethics

All studies using primary human lung material were approved by Hamilton Integrated Research Ethics Board (5099-T).

#### Primary Human Airway Epithelial Cells

Primary human bronchial epithelial cells (HBECs) were isolated immediately from bronchial brushing into T25 flasks containing Pneumacult™ Ex-Plus Basal Media (Stemcell Technologies, Vancouver Canada) with Pneumacult™ Ex-Plus 50x Supplement, 0.01% hydrocortisone stock solution and 1% antibiotic-antimycotic. Once cultures achieved ~80% confluence, cells were passaged to a T75 flask. Cells were fed with Pneumacult™ Ex-Plus Basal Media on a two-day feeding cycle. Once cultures reached 100% confluency, cells were trypsinized and seeded into microfluidic devices. A syringe pump was used to feed cells cultured in microfluidic devices every day. The feeding flow rate was 200 μL min^−1^, the devices were fed for 5 minutes, and the supernatant was collected for further possible analysis.

Calcein AM (5μM, Life Technologies®) dye was used to conduct a quantitative viability assay on day 7. First, the medium was removed from all microfluidic devices, at the cells were gently rinsed with warmed PBS using a 10-mL syringe, 400 μL of Calcein AM was added to each microfluidic device, they were incubated with Calcein AM dye at 37 °C for 20 minutes, the dye was removed, the microfluidic devices were rinsed with warmed PBS using a 10-mL syringe, and they were imaged by an EVOS M7000 microscope (Thermo Fisher, Canada).

### Coating Optimization

In this work, we developed and optimized PDA coating for microfluidic devices (Scheme 1a and Scheme 1b), where the coating was used as a linker to bind collagen inside the device. The height of the microfluidic channel was designed to be tall enough (~ 550 μm) to allow for further analysis of coated surfaces by cutting the channel and exposing the inner surfaces. Two methods were developed to coat PDA in microfluidic devices: 1) a simple one-step process in which DA solution was injected through the device and stored at room temperature for 24 or 48 hours (named as a static coating) and 2) a dynamic coating in which DA solution was continuously perfused through microfluidic devices in a closed-loop as shown in Scheme 1c. Scheme 1d also represents the coating process and conditions for PDA and collagen coating.

**Scheme 1:**
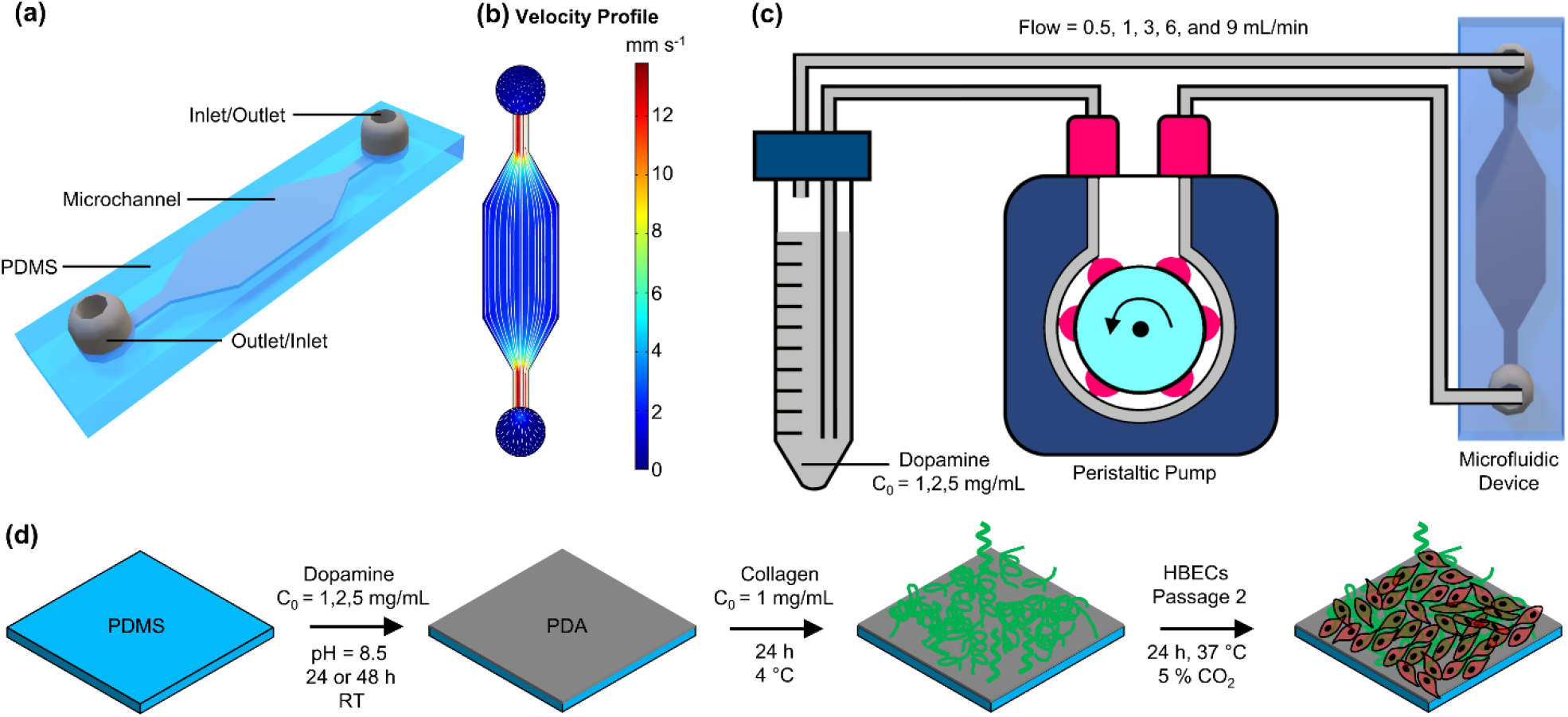
Representation of microfluidic device and the coating process: (a) 3D schematic view of the microfluidic device, (b) the velocity profile for the microfluidic device at a flow rate of 0.5 mL min^−1^ with flow streamlines showing the uniformity of flow inside of the device, (c) the PDA dynamic coating process using a peristaltic pump, and (d) the coating process of PDA and collagen.

## Results and Discussion

In the beginning, the effect of flow rate, DA concentration, and duration of the coating was investigated by measuring the water contact angle of the PDMS-coated devices. The aim of the characterization step was to optimize the coating recipe for a microfluidic device. In the first step, the impact of coating time under a constant DA concentration of 2 mg mL^−1^ was studied (Figure 1a and b). The water contact angle for the top side and the bottom side was separately measured to ensure that the coating was uniform and resulted in similar hydrophilicity. Generally, the water contact angle was varied from ~ 50° to ~ 70°, and the measurements among various conditions were not significantly different. However, an increase in coating time led to larger variation between the top and bottom sides of the channels showing that some non-uniformity or aggregation of nanoparticles could occur^18^. Figure 1c displays the average water contact angle for both 24 hours and 48 hours of PDA coating. The water contact angle was gently raised by increasing the flow rate but not significantly. However, the data was distributed closely around the average water contact angle when the coating time was 24 hours, and the flow rate of 0.5 mL min^−1^ or 1 mL min^−1^ for dynamic coating. Figure 1d shows the average water contact angle for static and dynamic coating (flow rate = 0.5 mL min^−1^ and 1 mL min^−1^) after 24 hours coating with three different DA concentrations. Higher DA concentration led to a small increase in the average water contact angle, especially for the static condition, and a larger variation in the data distribution was observed. Therefore, the DA concentration was chosen to be 2 mg mL^−1^ for the rest of the study, and the flow rate was fixed at 0.5 mL min^−1^ for dynamic coating. Next, the stability of PDA coating (for both static and dynamic conditions) was assessed by connecting the coated devices to a closed-loop fluid circuit in which deionized water (DI) continuously flowed at a flow rate of 10 mL min^−1^. The average water contact angle was measured at day 0, day 1, day 3, day 7, and day 14 without showing any noticeable change suggesting that the PDA coating was stable (Figure 1e). The roughness and topography of various treated and non-treated PDMS surfaces with PDA were also analyzed using AFM. Figure 1f depicts the root-mean-square (RMS) roughness of non-treated PDMS surfaces (native) and treated PDMS surfaces with PDA coating using the static and dynamic coating methods. The RMS roughness of native PDMS surfaces was 2.108 nm, which can be considered a regular measurement for PDMS. Nonetheless, the RMS roughness of PDA-coated PDMS surfaces was increased to 14.267 nm and 18.375 nm for the dynamic and static methods, respectively. This probably occurred because of the aggregation of PDA molecules on the PDMS surface, which could result in the formation of PDA nanoparticles^19,20^. Surface topography of the non-treated PDMS surface and the treated PDMS surfaces by PDA coating (static and dynamic) is presented in Figure 1g to Figure 1i.

**Figure 1:**
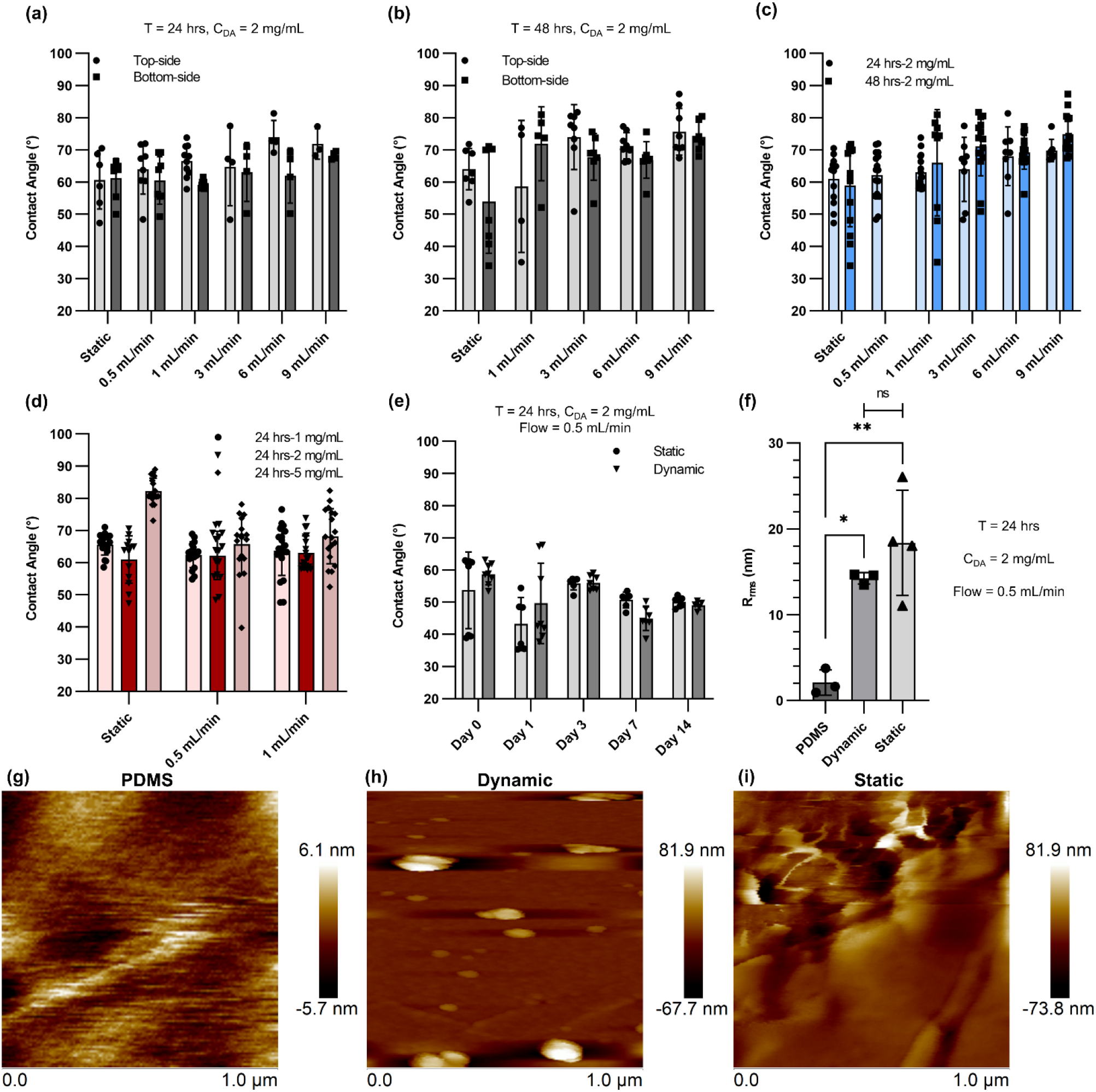
Characterization of PDA for microfluidic device: (a) contact angle data for static PDA coating and dynamic PDA coating with different flow rates after 24 hours coating with a concentration of 2 mg mL-1, (b) contact angle data for static PDA coating and dynamic PDA coating with different flow rates after 48 hours coating with a concentration of 2 mg mL-1, (c) comparison of contact angle data for 24 hours and 48 hours coating of PDA, (d) the effect of DA concentration on contact angle measurement for static and dynamic coating, (e) contact angle data for static PDA coating and dynamic PDA coating (flow rate = 0.5 mL min-1) over 14 days tested under a continuous flow, (f) surface roughness comparison of various conditions measured by AFM, (g) surface topography of a native PDMS surface, (h) surface topography of a treated PDMS surface coated with PDA using dynamic coating technique (flow rate = 0.5 mL min-1), and (i) surface topography of a treated PDMS surface coated with PDA using static coating method.

Primary human bronchial epithelial cells (HBECs) were cultured (cell density was 1.5 × 10^5^ per device) into untreated and treated devices to evaluate the biocompatibility of various coating conditions compared to native PDMS as seen in Figure 2a. Moreover, the covered surface area of adherent HBECs were quantified as a percentage of the total surface area using ImageJ, as shown in Figure 2b. HBECs on non-treated PDMS devices exhibited the lowest attachment (~ 2%) and the lowest growth over five days, confirming that native PDMS had poor surface biocompatibility for HBECs. In contrast, the cells on treated PDMS devices coated physically with collagen displayed a higher attachment and density compared to those on native PDMS surfaces. However, the adherent cell area for collagen treated PDMS devices decreased from day 1 to day 5, showing lifting of cells. The cells adhered at a higher density to the other two collagen-PDA-coated devices, and they could slowly spread over five days. On day 1 and day 2, the adherent cell density between the two conditions (PDMS+PDA-Static+Collagen and PDMS+PDA-Dynamic+Collagen) was not significantly different. At day 4 and day 5, a discernible difference between the adherent cell density was observed, suggesting that PDA-dynamic coating could provide a more suitable surface for cell proliferation. This observation is supported by AFM results. The static PDA coating could lead to more formation of PDA nanoparticles compared to the dynamic PDA coating, thereby increasing the surface roughness.

**Figure 2:**
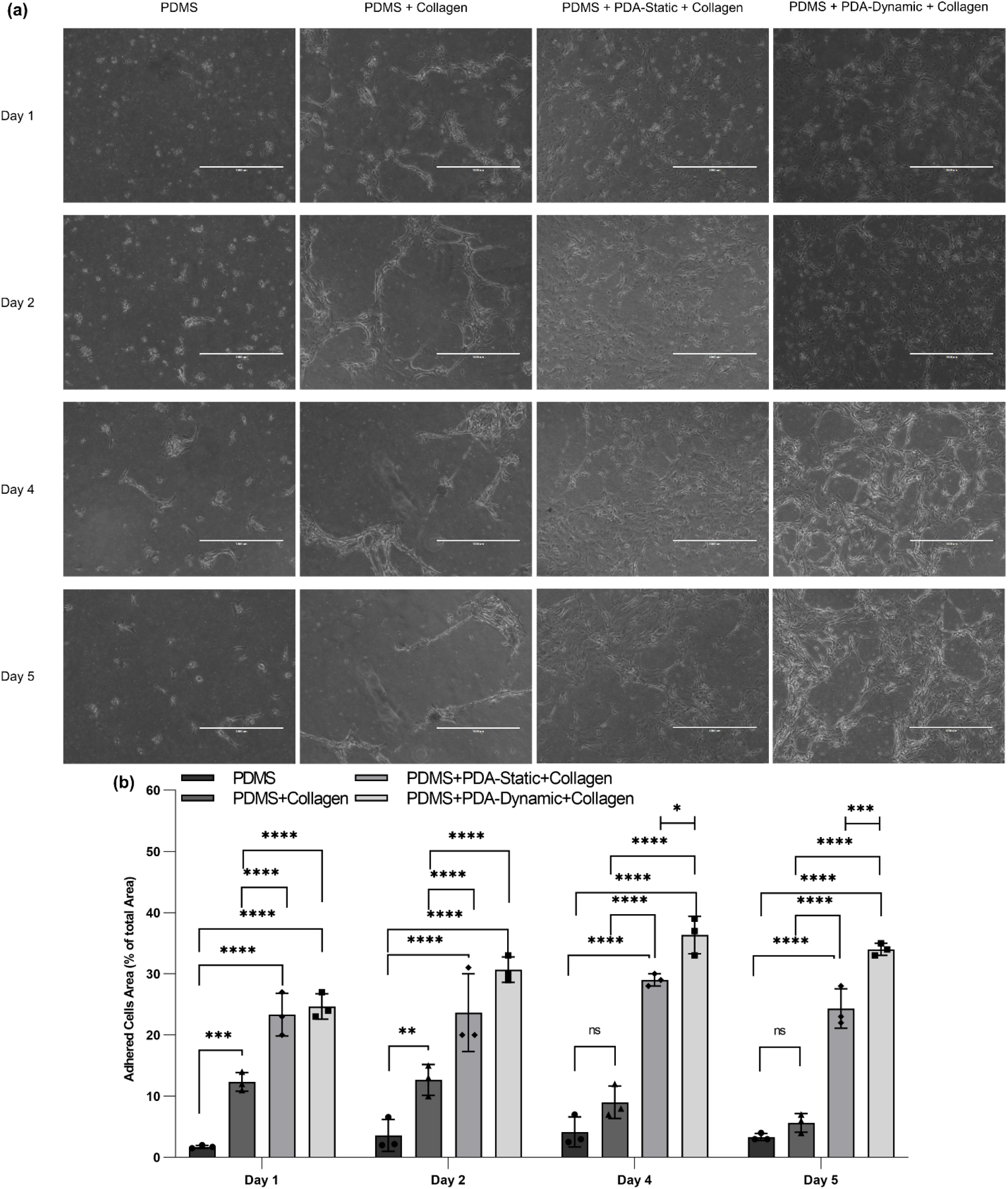
The comparison of cultured human bronchial epithelial cells (HBECs) on PDMS and treated surfaces: (a) bright-field image of HBECs adhered on native, collagen-coated, PDA-static-collagen-coated, and PDA-dynamic-collagen-coated surfaces at day 1, 2, 4, and 5 and (b) the adhered cells area as a percentage of the total area using bright field images that were quantified by ImageJ software. Scale bars are 1000 μm.

The cells were challenged by a flow stress test where the cells were exposed to a high flow rate for one minute. Such a sudden increase in shear stress was used to challenge the adhesion of cells to the surface. Moreover, a weaker collagen bonding would be prone to degradation, thereby detaching cells from the surface. Then, their viability and interleukin 8 (IL-8) (a marker of inflammation routinely studied in epithelial cell experiments^21–23^) were measured before and after the stress to explore the effect of the PDA coating method on cell viability and functionality. Since the cells did not adhere properly to native PDMS devices and collagen-coated PDMS devices (without PDA coating), these two groups were excluded from further studies. The next aim of the work was to compare the static and dynamic PDA coating using the flow stress test. The production level of IL-8 on day 1 and day 5 (before the flow stress test) was measured, as shown in Figure 3a. No perceptible difference between the two conditions was observed. Nevertheless, the production of IL-8 for devices with the dynamic PDA coating on day 5 was significantly higher than day 1. The cell density for the dynamic PDA-coated samples was increased by 40 % from day 1 to day 5, while the cell density for the static PDA-coated samples only exhibited 5% growth. As a result, the increase in IL-8 production could occur because there were more viable cells on day 5 for the dynamic PDA-coated samples compared to day 1. For the flow stress test, the flow rate was increased to 10 mL min^−1^ for one minute, and the supernatant was collected for further measurement, after feeding the devices at a normal flow rate of 200 μL min^−1^. Optical microscopy was performed to examine the samples (Figure 3b and Figure 3c), confirming that the cells still adhered to the surface. Figure 3d exhibits the adhered cells density before and after flow stress for both static and dynamic PDA-coated devices without any significant change. Moreover, the production of IL-8 cytokine was evaluated after the flow stress test to examine the effect of the coating type on the inflammatory response of the cells to stress (Figure 1e). No significant difference between two PDA coating was observed, suggesting that the type of coating did not play a role in the production level of the inflammatory cytokine of IL-8 during the flow stress test.

**Figure 3:**
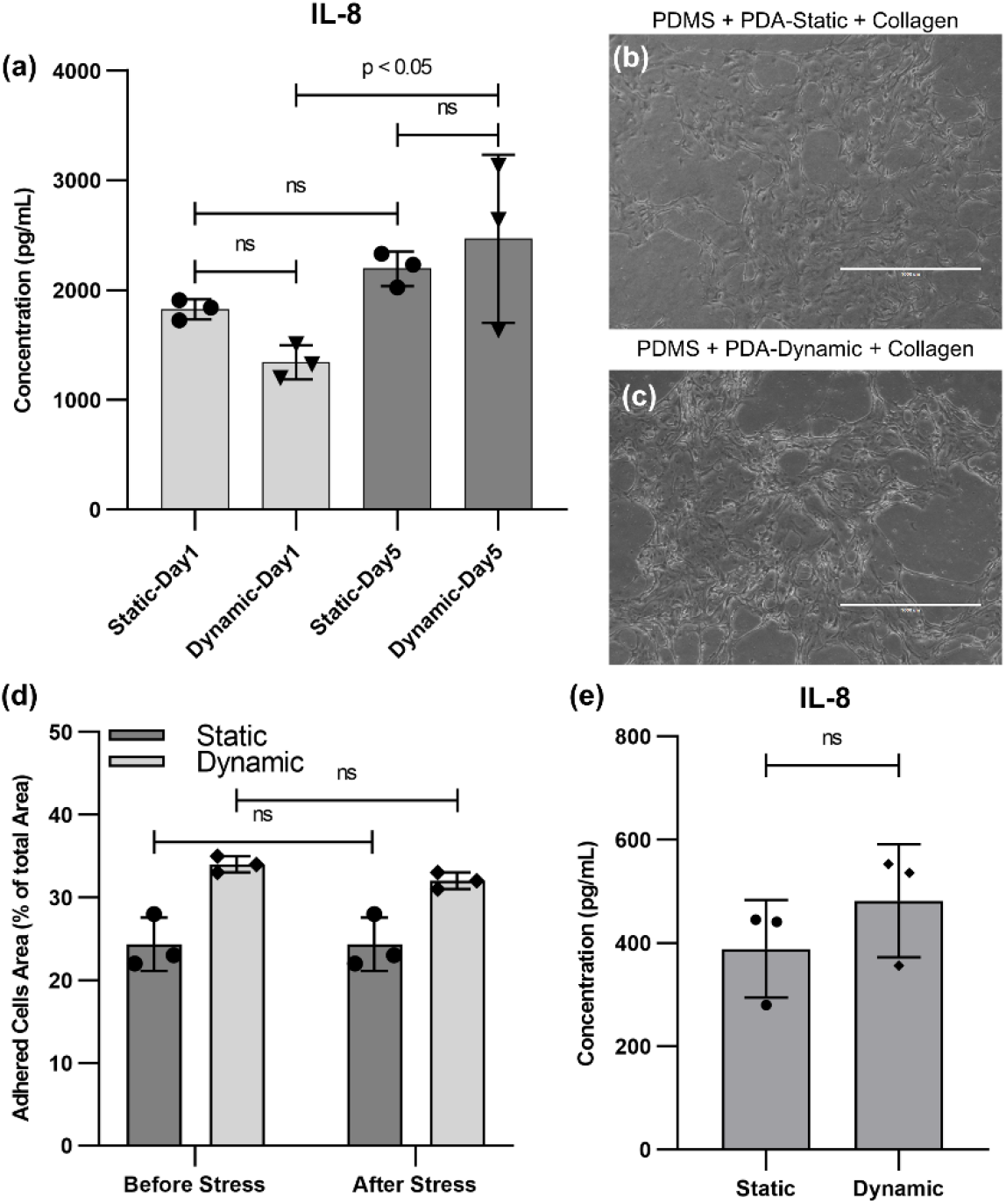
HBECs conditions before and after flow stress test: (a) IL-8 cytokine production by HBECs before flow stress at day 1 and day 5 for PDMS treated surfaces with collagen after dynamic or static PDA coating, (b) a bright filed image of HBECs on a PDMS treated surface with collagen using static PDA coating after the flow stress test, (c) a bright filed image of HBECs on a PDMS treated surface with collagen using dynamic PDA coating after the flow stress test, (d) a comparison of the adhered cells area as a percentage of the total area before and after flow stress testing for both dynamic and static PDA coating, and (e)) IL-8 cytokine production by HBECs after flow stress for PDMS treated surfaces with collagen after the dynamic and static PDA coating.

After the flow stress test, devices with PDA and collagen-coating were maintained for two additional days to ensure that the cells adhered to the surface as displayed in Figure 4a. For both static and dynamic coating, the cell density did not change after the flow stress test (Figure 4b), and no significant difference between the static and dynamic coating was observed. At day 7, the samples were stained with Calcein AM and imaged, as seen in Figure 4c and Figure 4d, demonstrating that the cells were alive and were not impacted by the flow stress test. Moreover, the cell density at the inlet and outlet of devices, where velocity and shear stress were higher, was lower.

**Figure 4:**
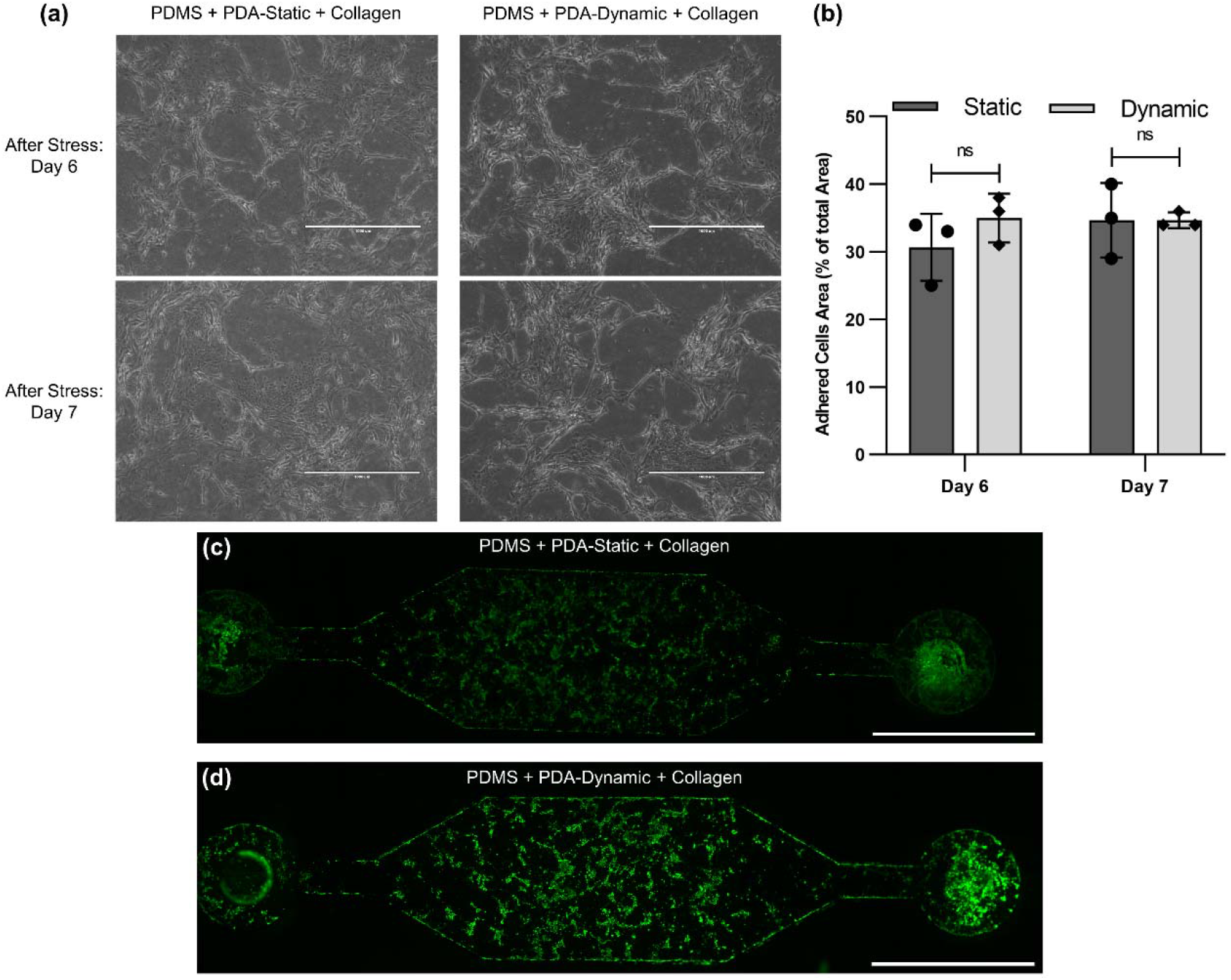
HBECs after flow stress test: (a) bright-field image of HBECs adhered on PDA-static-collagen-coated and PDA-dynamic-collagen-coated surfaces at day 6 (one day after flow stress test) and day 7 (two days after flow stress test), (b) the adhered cells area as a percentage of the total area at day 6 and day 7 for static and dynamic conditions, and a fluorescence image of HBECs adhered in a PDMS treated microfluidic device using (c) PDA-static-collagen-coating and (d) PDA-dynamic-collagen-coating.

## Conclusion

In summary, this work presented and compared a static and dynamic PDA coating for microfluidic devices. Different parameters of these static and dynamic PDA coating were studied and optimized by measuring the water contact angle and obtaining AFM images. PDA coating was used to bind collagen inside the microfluidic devices to improve the cell adhesion of primary human bronchial epithelial cells. Cell density, cytokine production, and resistance to a flow stress test were used to compare the effect of the coating method on cells. Cells could adhere properly to PDA-collagen-coated surfaces using both coating methods, and no significant difference between these two methods was observed. These results suggested that either the static or dynamic PDA coating can be applied to tailor PDMS-based microfluidic devices for binding an ECM protein. Such surface modification can have an application in micro-scale cell culture systems or organ-on-a-chip devices that stable long-term surface modification is sought.

## Acknowledgment

Dr. Jeremy Hirota would like to acknowledge the support from the SickKids New Investigator Award program and the Canada Research Chair in Respiratory Mucosal Immunology.

## Data Availability

The data that support the findings of this study are available from the corresponding author upon reasonable request.

